# Deep-tissue mechanosensation emerges from the interaction of external force and internal tissue state

**DOI:** 10.64898/2026.08.11.744238

**Authors:** Maximilian Nagel, Jonathan T. Seaman, Lars J. von Buchholtz, Nathan Ashby, Sienna K. Perry, Caroline Pierotti, Elizabeth A. Ronan, Joshua J. Emrick, Alexander T. Chesler

**Affiliations:** National Center for Complementary and Integrative Health (NCCIH), National Institutes of Health, Bethesda, MD 20892, USA; National Institute of Dental and Craniofacial Research (NIDCR), National Institutes of Health, Bethesda, MD 20892, USA; Department of Biologic and Materials Sciences, University of Michigan, Ann Arbor, MI 48109, USA; National Institute of Neurological Disorders and Stroke (NINDS), National Institutes of Health, Bethesda, MD 20892, USA; Research Management, Vertex Pharmaceuticals Incorporated, 50 Northern Avenue, Boston, MA 02210, USA

## Abstract

Muscle sensation is often considered in terms of proprioception, but muscle pain illustrates that other types of sensory neurons are also involved. Here, we show that trigeminal neurons innervating the masseter muscle fall into three classes: Aβ low-threshold mechanoreceptors, Aδ high-threshold mechanoreceptors (Aδ-HTMRs), and peptidergic neurons (PEP). All three types are recruited by mechanical stimulation, with massage being a particularly effective stimulus. Chemogenetic activation of masseter Aδ-HTMRs and PEP neurons results in pain-like symptoms. Moreover, acute and chronic inflammation sensitize nociceptors, altering behavioral tolerance in an animal model of massage. Taken together, our results provide a framework for understanding muscle somatosensation and how massage of sore muscles may be painful yet beneficial.

**Significance:** Muscle sensation is usually most prominent when something goes wrong: after overuse, injury, or inflammation, even ordinary pressure or movement can become painful. Yet how mechanical force is detected in muscle, and when a stimulus becomes painful, remain poorly understood. Here, we identify a simple cellular organization for muscle mechanosensation in which three sensory-neuron classes encode force through graded population recruitment. Massage effectively recruited all three classes, including the two nociceptive populations. We further show that these nociceptive neurons are sensitized by acute and chronic inflammation, leading to enhanced recruitment during massage. These findings show that the representation of mechanical force in muscle can be shaped by tissue state and provide a foundation for understanding muscle soreness, pain, and therapeutic touch.

## Introduction

Every year, millions of people resort to massage to alleviate muscle pain and soreness (1). However, despite its widespread use, the physiological basis underpinning the relief provided by massage remains unclear. The somatosensory system detects an array of stimuli through innervation of skin and deep tissue, including muscles (2, 3). We are usually oblivious to much of this sensory input — for example, the mechanical stimulation of our clothing when we bend our arms does not register unless something is wrong (4). Similarly, proprioception allows subconscious and normally effortless coordination of muscles, limbs, and body posture (5). Yet in deep tissues, conditions such as arthritis, joint inflammation, or muscle injury can dramatically affect both normal movement and perception, with even small movements evoking intense pain (6–8). Muscle pain can also arise through overuse, and massage can be painful even when it provides relief (3, 9).

Recent advances in our ability to distinguish, map, and measure the activity of sensory neurons have provided a new understanding of how the somatosensory system normally detects and distinguishes a range of stimuli (2) and how this representation can be transformed to trigger pain (10, 11). Here we combined an array of anatomical and functional imaging techniques with animal behavior to explore the sensory innervation of muscle and how these neurons respond to stimulation, including massage, at baseline and during inflammation. We show that most neurons innervating muscle belong to transcriptomic classes commonly thought to be involved in eliciting pain. Notably, massage was a particularly effective stimulus for these neurons, and our data reveal how the peripheral representation of this popular therapeutic approach is changed during inflammation.

## Results

### Retrogradely labeled masseter afferents fall into three classes and are broadly mechanosensitive

To target muscle-innervating neurons, we injected AAV6.2-tdTomato into the masseter, the muscle that controls jaw closing and generates bite force. Whole-head clearing and light-sheet imaging revealed axon labeling in the mandibular nerve selectively targeting the site of injection. Tracing these connections to the trigeminal ganglion (TG) revealed a discrete cluster of labeled cell bodies in the ipsilateral mandibular division that projected centrally to the spinal trigeminal subnucleus caudalis (Sp5c) and upper cervical segments (Fig. 1A–B; fig. S1A–C). Motor neurons targeting the masseter were also prominently labeled using this approach. To better understand the nature of the sensory innervation, we next used multigene in situ hybridization (ISH) to classify TG neurons into nine transcriptomically distinct groups (12, 13; Fig. 1C; fig. S1D). Three classes accounted for 95% of the virally labeled neurons. The two major populations, peptidergic neurons (PEP; ∼50%) and Aδ high-threshold mechanoreceptors (Aδ-HTMRs; ∼30%), are generally thought of as nociceptors, express *Scn10a*/NAV1.8 and *Calca*/CGRP, and are responsive to high-intensity mechanical stimuli (10, 11). The remaining cells were largely classified as Aβ low-threshold mechanoreceptors (Aβ-LTMRs; ∼15%; Fig. 1D), but not as proprioceptors, whose cell bodies are located in the mesencephalic nucleus rather than the TG (14). Importantly, there was almost no viral labeling of skin-innervating NP1–NP3, Aδ-LTMR, or C-LTMR neurons that express *Fxyd2* and/or *Tmem233*.

**Fig. 1.**
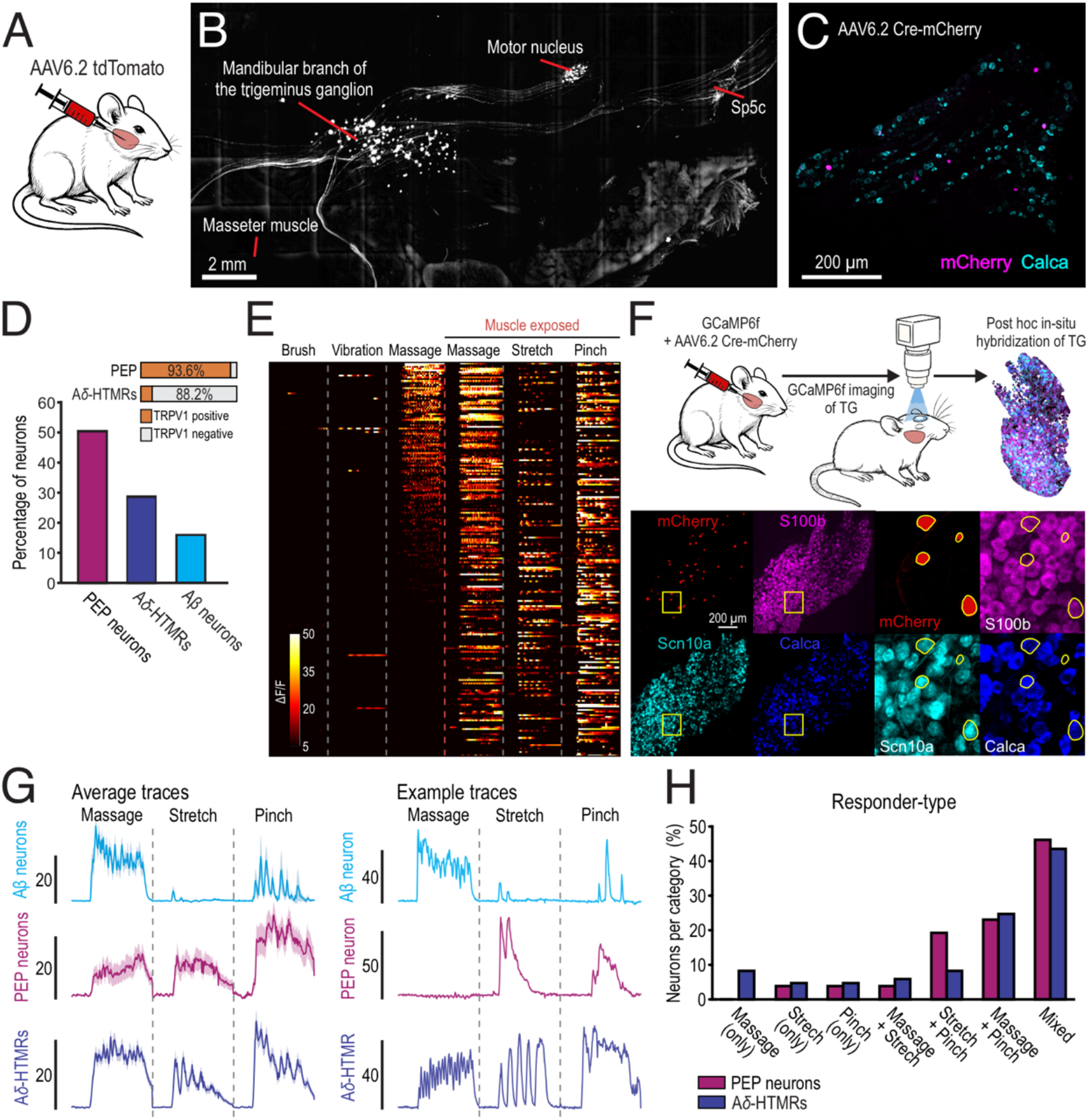
Masseter afferents are molecularly restricted and broadly responsive to deep-tissue force. (A and B) Whole-head tissue clearing and light-sheet imaging of AAV6.2-tdTomato labeling from the masseter revealed labeled axons in the mandibular nerve, somata in the mandibular division of the TG, and central projections to the spinal trigeminal subnucleus caudalis and C1–C2. Additional labeling was observed in the motor nucleus, whereas labeled neurons in the mesencephalic trigeminal nucleus, if present, were too sparse to detect (n = 1 mouse). (C) Representative TG section showing mCherry^+^ masseter afferents and multiplexed in situ hybridization for Calca and mCherry. (D) Molecular classification of labeled masseter afferents. Most labeled sensory neurons were PEP neurons (50.3%, 313/622 cells) or Aδ-HTMRs (28.6%, 178/622), with a smaller Aβ population (15.9%, 99/622); NP1 neurons were rare (0.3%, 2/622; not shown). Trpv1 expression was largely confined to PEP neurons. n = 622 neurons from 4 ganglia and 3 mice. (E) Population heatmaps showing mechanically evoked responses in TG neurons after AAV6.2-Cre-mCherry injection into the masseter muscle of GCaMP6f mice. Brush, vibration, and massage were first applied to the intact cheek. After removal of the overlying skin, massage, stretch, and pinch were applied directly to the exposed masseter. Each stimulus segment shows 40 s. GCaMP6f^+^ neurons were largely unresponsive to brush and vibration of the cheek, whereas cheek massage and direct stimulation of the exposed masseter recruited overlapping myofascial neurons. n = 224 neurons from 5 mice. (F) Strategy for post hoc molecular classification of functionally imaged myofascial sensory neurons. Representative whole-mount TG showing imaged neurons and post hoc in situ hybridization for mCherry, S100b, Scn10a, and Calca. (G) Average and example traces of molecularly identified Aβ, PEP, and Aδ-HTMR neurons showing mechanically evoked responses across stimulus modalities. Average traces are shown for all neurons within each class ± s.e.m. n = 7 Aβ neurons, 26 PEP neurons, and 85 Aδ-HTMR neurons from 5 mice. (H) Quantification of response profiles for PEP and Aδ-HTMR populations shown in (G). Both subtypes showed overlapping responses to massage, stretch, and pinch.

We next asked which stimuli activate these myofascial, muscle-innervating neurons by targeting GCaMP6f to the masseter using the retrograde viral approach (AAV6.2-Cre into Ai95 mice). As expected, labeled neurons were mostly unresponsive to gentle stimuli applied to the skin, including brush and vibration (Fig. 1E). By contrast, manual massage-like stimulation reliably elicited responses from myofascial neurons. Since GCaMP-expressing neurons were retrogradely labeled from the masseter, we predicted that their responses would be preserved after removal of the overlying skin. Indeed, directly massaging the exposed muscle elicited robust Ca^2+^ transients both from the neurons that were responsive through the skin and from additional cells (Fig. 1E). Other mechanical stimuli, including stretch induced by jaw movement and muscle pinch, were also effective at activating virally labeled neurons. To determine whether the three classes of muscle-innervating neurons were differentially tuned, we carried out *post hoc* ISH (15; Fig. 1F). Notably, while PEP neurons and Aδ-HTMRs were broadly responsive to all stimuli, the smaller number of Aβ neurons responded primarily to massage (Fig. 1G–H, fig. S2).

### Myofascial sensory neurons encode mechanical force largely independently of Piezo2

As an alternative approach to broadly target masseter-innervating neurons, we used a strategy of neonatal AAV9-Cre injection into Ai95 mice to drive stochastic GCaMP6f expression across all classes of TG neurons (16). We reasoned that, in this preparation, many neurons would be responsive to sensory stimuli, but myofascial neurons would continue to respond during muscle stimulation after removal of the overlying skin. As expected, many LTMRs and HTMRs were recruited in response to brush, vibration, and pinch applied to the intact cheek (fig. S3A). After removal of the skin, TG neurons rapidly became silent, with GCaMP fluorescence returning to baseline within minutes, suggesting that cutting peripheral projections does not induce prolonged firing or sustained Ca^2+^ elevation. A select subset of TG neurons remained responsive to muscle stimulation (Fig. 2C). To verify that these cells innervated the muscle rather than surrounding tissue, we focally applied 500 mM KCl to selectively depolarize terminals in the masseter. The KCl-responsive neurons exhibited tuning that closely recapitulated the activity profiles of retrogradely labeled masseter neurons (Fig. 1H).

**Fig. 2.**
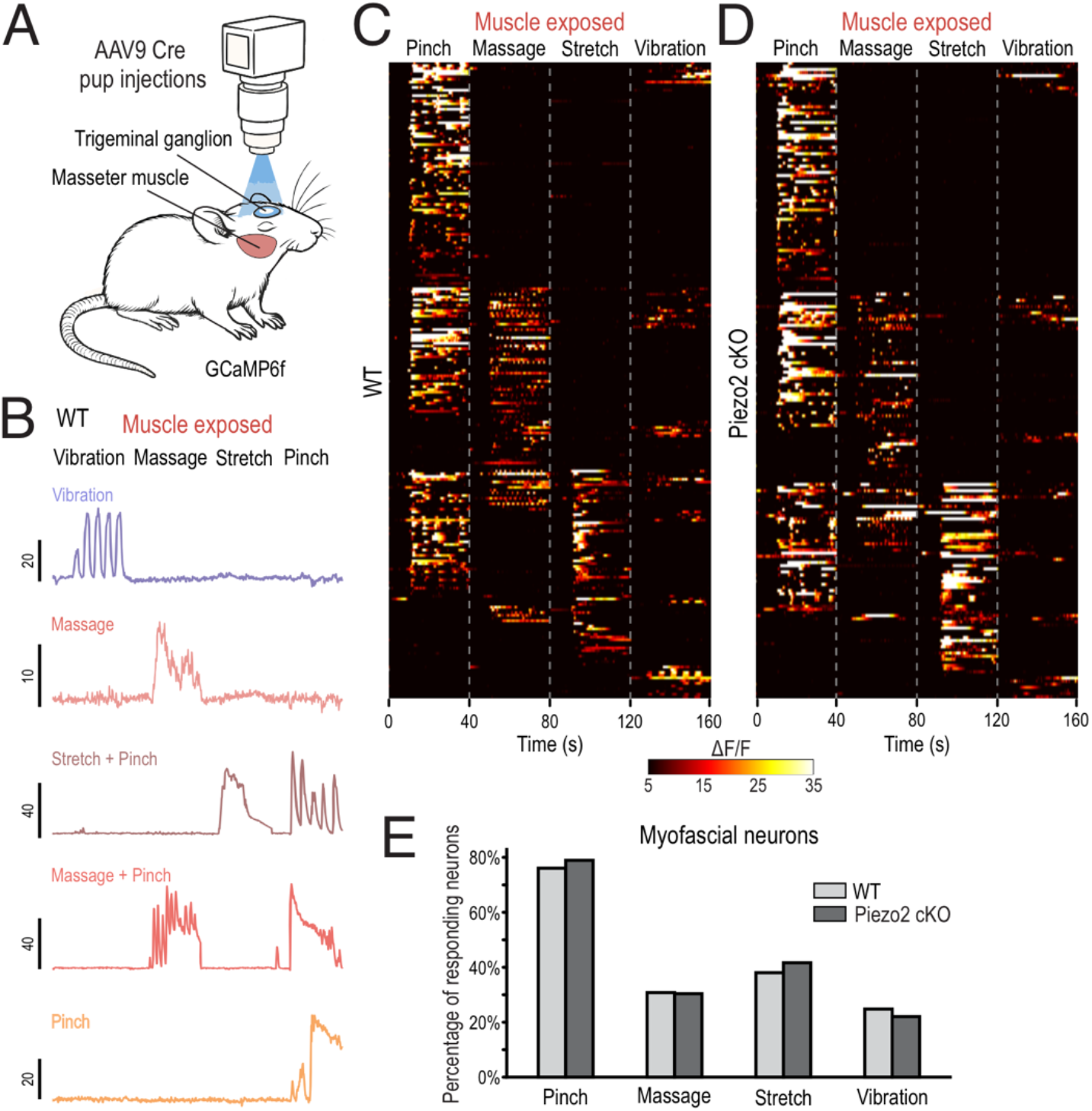
Myofascial sensory neurons encode mechanical stimuli largely independently of Piezo2. (A) Schematic of the in vivo Ca^2+^ imaging preparation enabling direct mechanical stimulation of the exposed masseter muscle while imaging TG neurons. Neurons responding to KCl application and direct mechanical stimulation of the exposed masseter were classified as myofascial afferents. (B) Representative Ca^2+^ responses of myofascial afferents to direct mechanical stimulation of the exposed muscle, including vibration, massage, stretch, and pinch. (C) Population heatmaps of mechanically evoked responses across stimulus modalities in WT mice. Pinch, massage, stretch, and vibration recruited overlapping subsets of myofascial neurons, with pinch recruiting the largest fraction of the population. Frequency-tuned responses to vibration applied directly to muscle were rare. n = 234 neurons from 3 mice. (D) Same analysis in Piezo2 conditional knockout mice. Responses to massage, stretch, and pinch were largely preserved, whereas vibration responses lost frequency dependence and were consistent with tissue displacement rather than frequency-tuned vibration responses. n = 204 neurons from 4 mice. (E) Quantification of mechanically evoked response classes in WT and Piezo2 conditional knockout mice. The percentage of responders to each stimulus did not differ significantly between genotypes (Fisher’s exact test with Holm correction: all adjusted p = 1.0).

The mechanotransduction channel Piezo2 is expressed in a subset of Aδ-HTMRs as well as Aβ-LTMRs and has been shown to be important for their functional responses (16–18). After Piezo2 knockout, skin neurons lost responses to brush and vibration (fig. S3B). By contrast, responses from muscle-innervating neurons were largely unaltered, including responses to massage, stretch, and pinch (Fig. 2D–E). This finding is fully concordant with our previous work on human subjects with PIEZO2 deficiency, who retain sensitivity to deep-tissue pressure despite their profound loss of proprioception and discriminative touch (19).

### Activation of *Scn10a*^+^ myofascial neurons induces orofacial pain

Aδ-HTMRs and PEP neurons innervating the skin function as nociceptors (20). These cell classes express *Scn10a* (17; Fig. 1D; fig. S1D), allowing us to target the two major populations innervating the masseter using *Scn10a*-Cre (21; Fig. 3A). Reporter labeling in these mice revealed a dense network of sensory axons throughout the masseter, forming a fine mesh of free nerve endings without obvious specialized terminal structures, muscle spindle endings, or motor neuron endings (Fig. 3B–D). To assess nocifensive responses evoked by activating *Scn10a*-expressing neurons, we used a chemogenetic strategy and focal CNO delivery. As expected, subcutaneous CNO injection into the hindpaw evoked classic nocifensive behaviors, including licking and shaking of the injected paw (fig. S4A–D), confirming chemogenetic activation. By contrast, low-volume CNO injection targeted to the masseter failed to evoke facially directed nocifensive behaviors. Instead, animals exhibited freeze-like behavior and a striking reduction in movement in an open-field assay (Fig. 3E–H), consistent with our previous studies of orofacial pain (22).

**Fig. 3.**
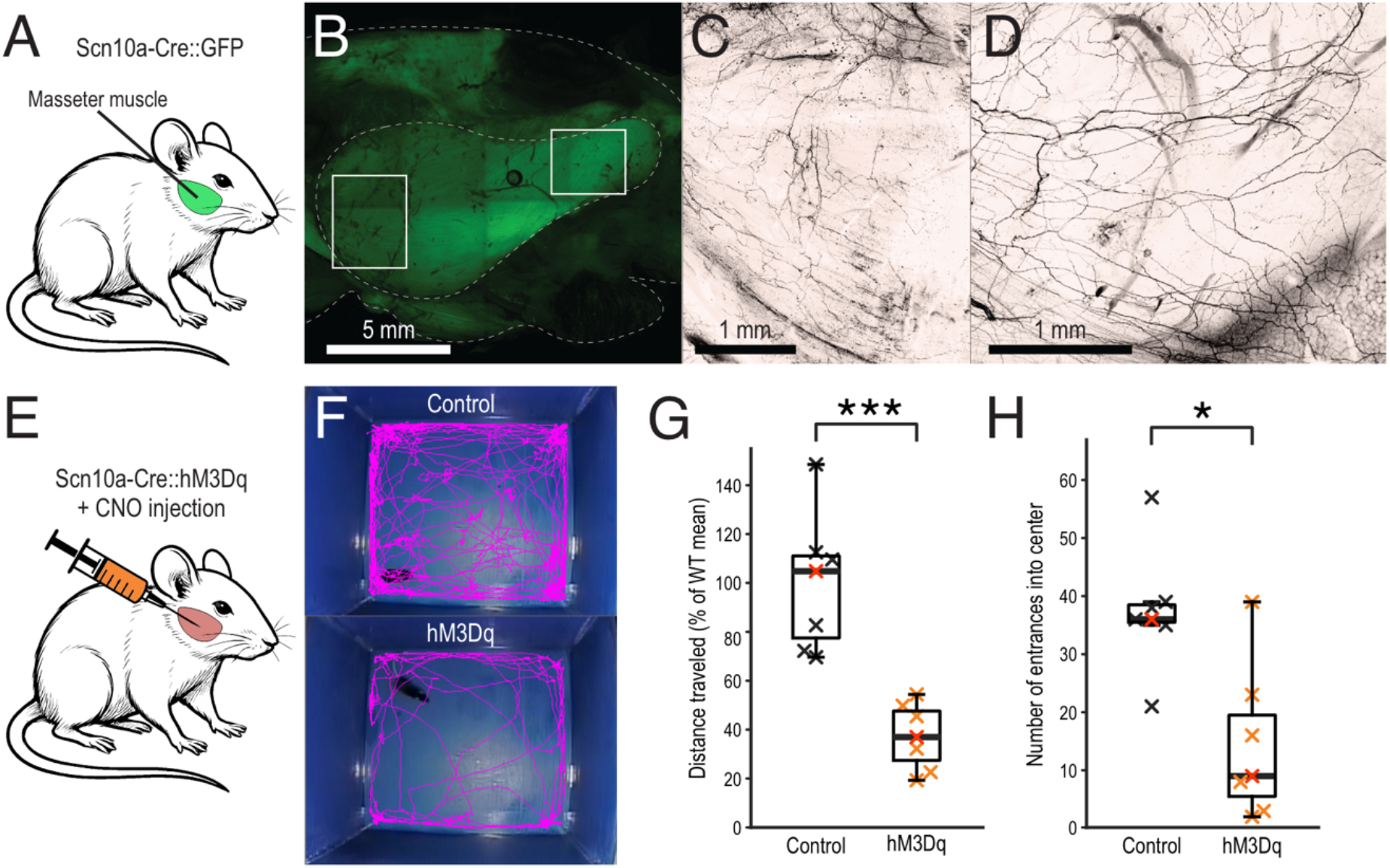
Scn10a^+^ myofascial afferents form free nerve endings and induce orofacial pain. (A) Scn10a-Cre mice were used to label a broader myofascial population, as approximately 80% of masseter afferents belonged to Scn10a-expressing nociceptor classes. (B to D) Reporter labeling revealed a dense network of GFP^+^ sensory axons throughout the masseter, forming a fine mesh of free endings without obvious specialized terminal structures, muscle spindle endings, or motor neuron endings. (B) Overview image of labeled axons in the masseter. (C) Higher-magnification view of the region outlined in the left box in (B). (D) Higher-magnification view of the region outlined in the right box in (B). n = 3 mice. (E) Schematic illustrating intramuscular injection of CNO into the masseter muscle of Scn10a-Cre::Rosa26-hM3Dq mice. WT and hM3Dq mice received identical CNO dosing. (F) Representative movement trajectories of WT and hM3Dq mice in an open-field arena after masseter CNO injection. (G) Distance traveled in the open-field arena, normalized to the WT mean. hM3Dq mice show a marked reduction in total movement compared to WT controls (WT: 100%; hM3Dq: 37.3%; Mann–Whitney U test, p = 0.0006; n = 7 mice per group). (H) Number of center entries in the open-field arena. hM3Dq mice exhibit significantly fewer center entries than WT controls (WT: 37.4; hM3Dq: 14.3 entries; Mann–Whitney U test, p = 0.025; n = 7 mice per group). Representative mice shown in (F) are indicated with a red X.

### Inflammation induces subtype-specific shifts in mechanical sensitivity

Classic studies by Mense and colleagues in cats have suggested that complete Freund’s adjuvant (CFA)-induced inflammation sensitizes muscle afferents to a variety of different types of stimulation (23). To better define how inflammation reshapes mechanosensory responses of nociceptors, we injected CFA into the masseter muscle of *Scn10a*-Cre::GCaMP6f mice and imaged TG responses to defined forces using von Frey filaments (Fig. 4A–B). PEP neurons express TRPV1 and respond to capsaicin, providing a simple method to distinguish them from Aδ-HTMRs by measuring their sensitivity to capsaicin (Fig. 1D; figs. S5 and S6). As before, we defined myofascial neurons by their responses to chemical stimulation delivered directly to the muscle (capsaicin, 500 mM KCl, or both; fig. S5). Without inflammation, both PEP neurons and Aδ-HTMRs showed progressive recruitment with increasing force across the tested range (Fig. 4A–B). In CFA-injected animals, PEP neurons were noticeably more sensitive across a wide range of mechanical stimulation (2–15 g; Fig. 4A–C; fig. S7A). However, Aδ-HTMRs showed no significant net change related to inflammation (Fig. 4A, fig. S7B). This subtype-specific effect of inflammation parallels prior *ex vivo* findings in incised muscle (24). We next examined how the von Frey-sensitive neurons responded to noxious stimulation. As expected, pinch activated almost all of these neurons both at baseline and after CFA injection (fig. S7C).

**Fig. 4.**
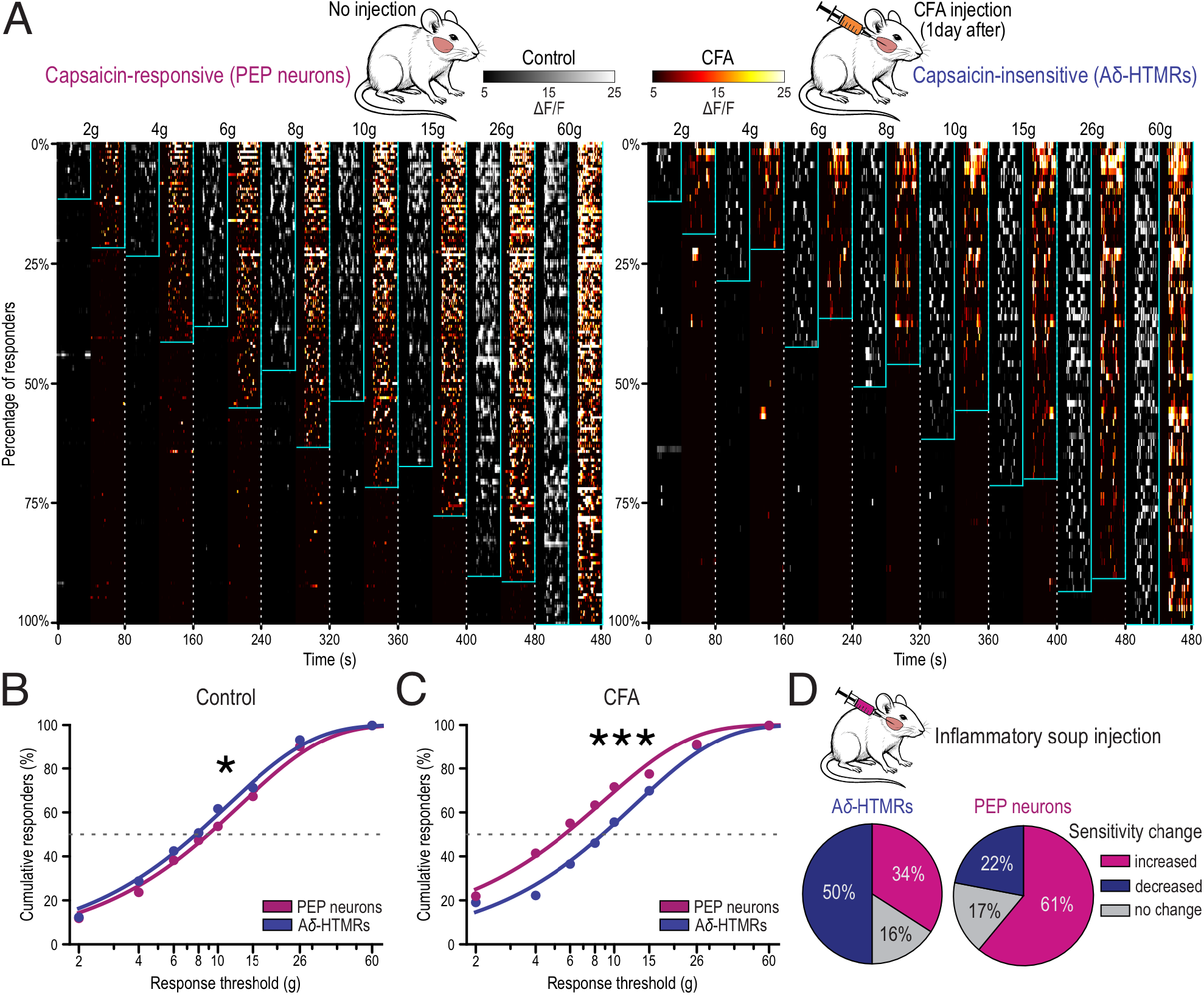
Inflammation shifts myofascial mechanoreceptor sensitivity in a subtype-specific manner. (A) Top, experimental setup for induction of chronic inflammation by intramuscular CFA injection into the masseter muscle of Scn10a-Cre::GCaMP6f mice, followed by in vivo Ca^2+^ imaging of TG neurons 24 h later. Left, population heatmaps show alternating von Frey responses (2–60 g) for capsaicin-responsive PEP neurons (gray, control; fire, CFA). Control: n = 110 neurons from 5 mice; CFA: n = 169 neurons from 6 mice. Right, same analysis for capsaicin-insensitive Aδ-HTMRs. Control: n = 73 neurons from 5 mice; CFA: n = 63 neurons from 6 mice. (B) Force–response curves under control conditions. Aδ-HTMRs were more mechanically sensitive than PEP neurons (ΔEC50 = 1.19 g; PEP EC50 = 8.95 ± 0.46 g; Aδ EC50 = 7.76 ± 0.46 g; nonlinear regression comparison, F-test p = 0.0238). (C) Force–response curves under CFA conditions. CFA increased mechanical sensitivity of PEP neurons, reversing the subtype relationship observed under control conditions (ΔEC50 = 2.56 g; Aδ EC50 = 8.74 ± 0.61 g; PEP EC50 = 6.18 ± 0.51 g; nonlinear regression comparison, F-test p = 3.45 × 10^-5^). (D) Longitudinal in vivo Ca^2+^ imaging during 2–10 g von Frey stimulation before and 4 min after acute inflammatory soup (IS) injection into the masseter muscle. Pie charts show changes in sensitivity following IS injection. Aδ-HTMRs showed heterogeneous changes in sensitivity (50% decreased, 34% increased, 16% no change), whereas the distribution of changes in PEP neurons was shifted toward increased sensitivity (61% increased, 22% decreased, 17% no change). Aδ-HTMRs: n = 44 neurons; PEP neurons: n = 100 from 8 mice.

In parallel, we took a different approach to sensitize the muscle by injecting a cocktail of inflammatory mediators (“inflammatory soup,” IS) that are commonly used to induce acute pain (25). This model allowed us to quantify changes in sensitivity of individual neurons as muscle inflammation develops (fig. S8A–B). Just as with CFA, the response properties of Aδ-HTMRs were largely unchanged after IS injection (Fig. 4D; fig. S8C, G). Again, PEP neurons were sensitized by IS injection, recapitulating the effect of CFA (Fig. 4D; fig. S8D, G).

### Inflammation enhances massage-evoked recruitment and reduces massage tolerance

We next asked whether this inflammatory model also impacts nociceptor recruitment during massage (Fig. 5A). Intriguingly, both PEP neurons and Aδ-HTMRs were sensitized to a similar extent following injection of IS. We also noted that these heightened massage-evoked responses decreased with repeated stimulation in both populations.

**Fig. 5.**
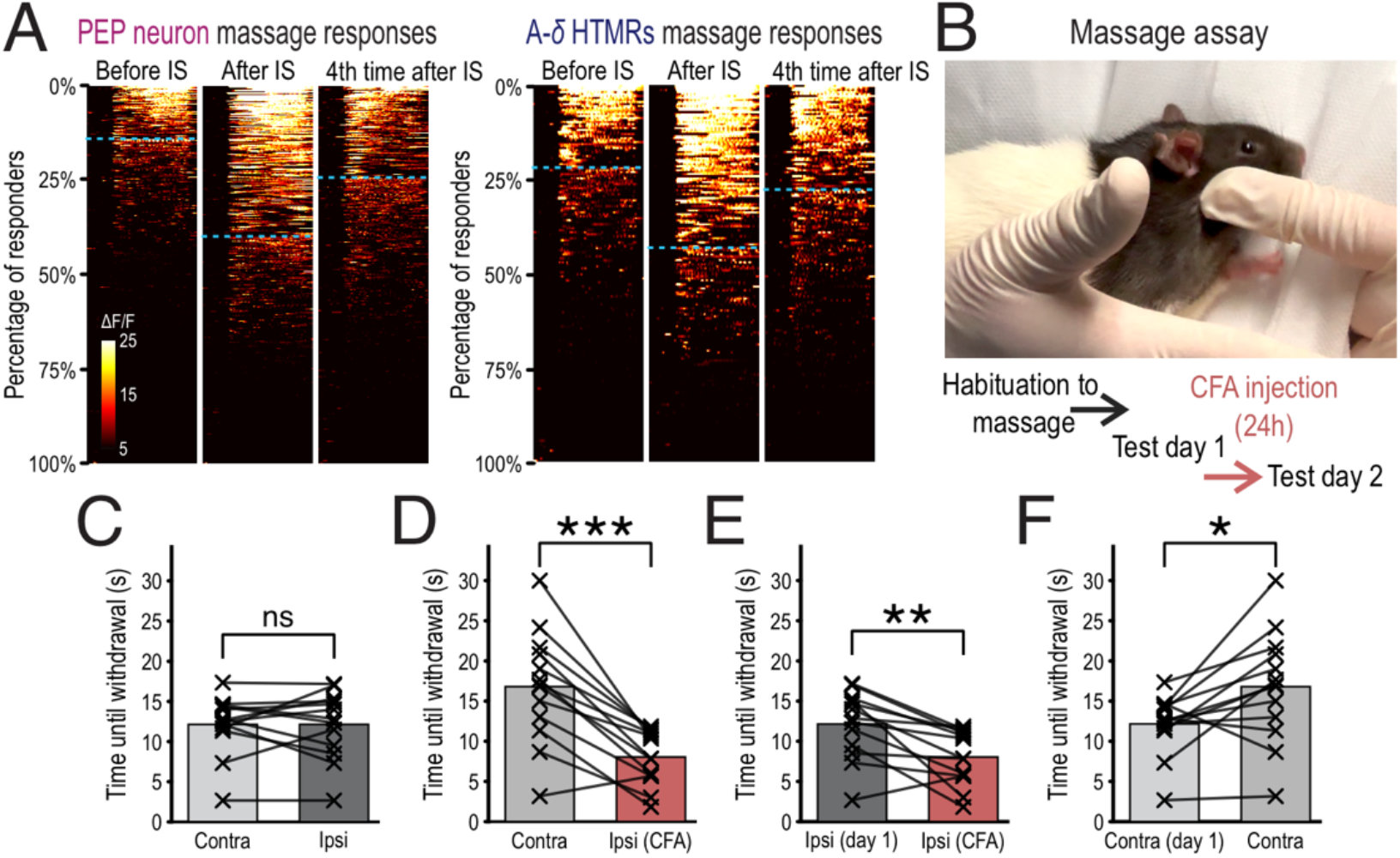
Inflammation reshapes massage-evoked recruitment and behavioral tolerance. (A) Scn10a-Cre::GCaMP6f Ca^2+^ imaging of massage-evoked responses in PEP neurons (left) and Aδ-HTMRs (right) before and after inflammatory soup (IS) injection into the masseter. Population heatmaps show sustained and transient responses, separated by a dotted line; sustained responses were defined as lasting at least 5 consecutive seconds. After repeated massage following IS, responses approached baseline levels by the fourth stimulation. Each stimulus segment was sorted independently and shows 40 s. Aδ-HTMRs: n = 188; PEP neurons: n = 468 from 8 mice. (B) Experimental design for assessing massage tolerance before and after unilateral CFA injection into the masseter muscle. Rats were habituated to the massage paradigm for 2 weeks. On test day 1 (baseline), each masseter was stimulated six times for up to 30 s, and withdrawal latency was measured. On test day 2, 24 h after unilateral CFA injection, withdrawal latency was reassessed on both sides. (C) Under baseline conditions, animals tolerated massage for comparable durations on both masseters (n = 13 rats; Wilcoxon signed-rank test, p = 0.97). (D) Following unilateral CFA injection, massage tolerance was significantly reduced on the inflamed side (p = 0.0005). (E) Withdrawal latency on the inflamed side was reduced relative to the same side at baseline (p = 0.0024). (F) Massage tolerance increased on the contralateral, non-inflamed side relative to baseline (p = 0.017).

Massage is a popular therapeutic approach used to soothe aching muscles. However, an effective massage can often be quite painful. For human subjects, this is particularly true when aching or injured muscle is massaged, although many people report pain starts to decrease over the course of the massage (26). This closely parallels the initial sensitization of nociceptors following IS injection and its subsequent decline with repeated massage stimulation (Fig. 5A). Mice are typically averse to human handling; therefore, we devised a rat model to test whether massage becomes less tolerable after inflammation. Rats were habituated to cheek massage and quickly learned to tolerate this type of manipulation, interacting with the investigator for extended periods before withdrawing (Fig. 5B–C; Video S1). By contrast, after inflammation, their behavior was substantially changed when massage was directed to the inflamed side (Video S2). This significantly reduced the interaction time for ipsilateral massage but modestly increased this measure when massage was applied to the unaffected cheek (Fig. 5D–F). Taken together, these animal studies reveal how massage activates muscle-innervating Aβ-LTMRs (Fig. 1), Aδ-HTMRs and PEP neurons (Figs. 1 and 5), and how the latter populations are sensitive to inflammation, closely matching the human experience of pain when massage is applied to inflamed tissue.

## Discussion

Deep tissue is richly innervated by the somatosensory system, yet we are rarely aware of sensations from our muscles during ordinary use, except following overuse or trauma (23). Here we show that just a small subset of neural classes target the masseter muscle of mice, with about 80% coming from classes typically thought of as nociceptors (Aδ-HTMRs and PEP neurons; 11) that respond to high-intensity mechanical stimuli. Chemogenetic activation of these nociceptor classes affected mouse behavior, and, although the changes we observed appeared relatively subtle, they were completely consistent with other trigeminal pain models (13, 22, 27, 28). Similarly, after the masseter was injected with CFA, we did not observe overt pain behaviors or changes in food intake or body weight, despite obvious swelling of the cheek. Nonetheless, both CFA and IS, two very different types of inflammatory agents, selectively sensitized PEP neurons to von Frey stimulation (Fig. 4), but more generally sensitized muscle-innervating HTMRs (Fig. 5). This provides a plausible cellular basis for the reduced tolerance to massage of sore muscles, both in humans (26) and in our rat model (Fig. 5).

How does nociceptor sensitization arise in response to inflammation? At the most basic level, it could reflect increased mechanosensitivity selectively in these classes of neurons (23). However, the increased mechanosensitivity would have to be a specialized property of masseter-innervating neurons, since no mechanical sensitization was observed during skin inflammation (13). Alternatively, sensitization could be a consequence of changes in the stiffness of the muscle affecting the force experienced by the afferent terminals. Yet both PEP and Aδ-HTMRs largely appear to form free nerve endings in the muscle (Fig. 3; 29), making it difficult to explain their differential properties in response to von Frey filaments during inflammation by changes in tissue compliance alone. It is also possible that, in response to mechanical stimulation, the damaged muscle changes the local environment, for example pH or ATP levels, and that PEP neurons are particularly sensitive to local changes induced by punctate force (30–32).

Our data establish how three classes of sensory neurons in the trigeminal ganglion are recruited in an overlapping, graded fashion by natural stretch and exogenous mechanical stimulation of the masseter. In addition, masseter proprioceptors, which are localized to the mesencephalic nucleus of the trigeminal in the brainstem, likely provide PIEZO2-dependent sensory input related to motor control (5, 14). A holistic understanding of how these separate mechanosensory pathways combine to produce a sensory percept and regulate motor function will ultimately require mapping the postsynaptic circuits in the brain (33). There is also the broader question of where the distinction between innocuous and noxious sensation arises, and how relatively subtle shifts in perceptual quality are related to the differential activity profiles we observed. Our results are compatible with a state-dependent shift in the relative sensitivity of muscle-innervating neurons as one driver of altered muscle sensation, ultimately providing a plausible foundation for understanding the relief afforded by massage of inflamed tissue.

## Acknowledgments

This study was supported in part by the Intramural Program of the National Institutes of Health, National Center for Complementary and Integrative Health and National Institute of Neurological Disorders and Stroke (A.T.C., ZIA AT000028), National Institute of Dental and Craniofacial Research (ZIA DE000561) and Department of Biologic and Materials Sciences, University of Michigan (J.J.E., UC2AR082197; E.A.R., T32 DE007057 and T32 DC00011; S.K.P., T32 DE007057). We thank Nicholas J. P. Ryba for constructive feedback and discussions during manuscript preparation. We thank members of our groups for input. ChatGPT-5 (OpenAI) was used to assist with refinement of manuscript text and with coding support for data analysis. All scientific content, code, analyses, outputs, and interpretation were reviewed and verified by the authors.

## Author contributions

M.N., J.J.E. and A.T.C. designed research; M.N., J.T.S., L.J.v.B., N.A., S.K.P., C.P. and E.A.R. performed research; M.N. and L.J.v.B. analyzed data; J.J.E. and A.T.C. supervised the work; and M.N., A.T.C. wrote the paper.

## Data and code availability

All data supporting the findings of this study are available from the corresponding author and will be deposited in a public repository prior to publication. Custom MATLAB and Python scripts used for data processing and analysis are available from the corresponding author upon reasonable request and will be publicly released upon publication.

## Supplementary Materials

Materials and Methods

Figs. S1 to S8

Video S1 and S2

