## Supplementary Materials for "Deep-tissue mechanosensation emerges from the interaction of external force and internal tissue state"

### Materials and Methods

#### Experimental animals

All animal experiments were performed in accordance with the guidelines of the National Institutes of Health and were approved by the Institutional Animal Care and Use Committee (IACUC) of the National Institute of Neurological Disorders and Stroke. The following mouse lines were obtained from The Jackson Laboratory: Ai95(RCL-GCaMP6f)-D (34; no. 024105), Nav1.8-Cre (21; no. 036564), RC::FL-hM3Dq (35; no. 026942), and Ai140(TIT2L-EGFP-ICL-tTA2) (36; no. 034100). The Piezo2$^{lox/lox}$ mouse line was described previously (16). Male and female mice were used in all experiments. Animals were housed under controlled environmental conditions (23 °C, 50% humidity, 12-h light/dark cycle) with ad libitum access to standard laboratory chow and water.

#### Viral vectors and genetic strategies

AAV serotype 9 (AAV9), an established serotype for neonatal transduction (16), was used for injections at postnatal day 1 (P1). For neonatal delivery, 10 µl of 1:10 diluted virus was injected subcutaneously close to the milk spot. AAV6.2 was chosen based on our previous use of this serotype to label tooth-innervating trigeminal sensory neurons (22, 37). Here, we established the use of intramuscular AAV6.2 injections in adult mice to transduce sensory neurons innervating the masseter muscle. Unless otherwise stated, AAV6.2 was used undiluted. The following viral constructs were used: AAV9-CAG-Cre ($2 \times 10^{12}$ vg/ml; Vigene; CV17187-AV9); AAV9-hSyn1-chI-mCherry_2A_iCre-WPRE-SV40p(A) ($5.6 \times 10^{12}$ vg/ml; Viral Vector Facility, Universität Zürich; p147); AAV6.2-hSyn1-chI-mCherry_2A_iCre-WPRE-SV40p(A) ($6 \times 10^{12}$ vg/ml; Viral Vector Facility, Universität Zürich; p147); AAV6.2-shortCAG-FLPo-WPRE-SV40p(A) ($1 \times 10^{12}$ vg/ml; Viral Vector Facility, Universität Zürich; p453); and AAV6.2-shortCAG-tdTomato-WPRE-SV40p(A) ($2 \times 10^{12}$ vg/ml; Viral Vector Facility, Universität Zürich; p131).

#### Surgical procedures, viral tracing, and induction of inflammation

To enable precise tracing and labeling of myofascial neurons innervating the masseter muscle, animals aged ≥8 weeks were anesthetized with isoflurane (2% in oxygen). A single skin incision was made over the masseter, and connective tissue was carefully separated to expose the muscle. Undiluted AAV6.2 was injected using a Hamilton syringe (30G needle) at five sites within the masseter (1.25 µl per site; total volume 6.25 µl), with two injections placed into the superficial masseter and three into the deep masseter. Injections were performed slowly, and the needle was left in place for 20–30 s before withdrawal to minimize reflux. To enable precise intramuscular injections through the skin to induce chronic inflammation, animals were briefly anesthetized with isoflurane. A single injection of complete Freund's adjuvant (CFA; 20 µl in mice, 100 µl in rats) was delivered into the muscle belly of the masseter muscle 24 h prior to experimentation. For acute inflammatory modulation during in vivo $Ca^{2+}$ imaging, inflammatory soup (IS) was injected into the exposed masseter muscle (10 µl into the superficial masseter and 10 µl into the deep masseter). IS consisted of serotonin (100 µM), bradykinin (100 µM), histamine (100 µM), and prostaglandin $E_2$ (10 µM), dissolved in PBS.

**In vivo calcium imaging of trigeminal ganglion neurons**

For fluorescent calcium imaging of trigeminal ganglion (TG) neurons, animals aged ≥8 weeks were anesthetized with isoflurane or ketamine and prepared for optical access to the TG as described previously, with imaging focused on the mandibular division (16). Ketamine anesthesia (80 mg/kg ketamine and 6 mg/kg xylazine) was used for experiments in *Scn10a*-Cre::GCaMP6f mice; isoflurane (1.5% in oxygen) was used for all other TG imaging experiments. The head was stabilized using custom anterior and posterior head bars to permit high-force mechanical stimulation without movement. Cutaneous stimulation was performed as indicated below. To access the masseter muscle, the cheek skin was removed by separating the connective tissue between skin and muscle. The exposed muscle was kept moist with PBS throughout the experiment. Although skin removal induced transient activation of cutaneous neurons, activity returned to baseline within 2–4 min before muscle stimulation commenced. Animals were subjected to one of three experimental regimes. Regime A (Fig. 1) employed a short stimulation protocol. After cutaneous stimulation (vibration, brushing, massage), massage, stretch, and pinch were directly applied to the exposed masseter. In these experiments, GCaMP6f expression was induced virally from the masseter muscle to increase specificity for myofascial afferents. Regime B (Fig. 2) assessed mechanical responses in both skin and muscle. Cutaneous stimuli included vibration, brushing, massage, jaw-induced stretch, and pinch. Muscle stimuli were delivered in increasing intensity (vibration, massage, jaw-induced stretch, pinch). At the end of each experiment, 0.5 M KCl was applied to the muscle surface and injected intramuscularly (20 µl into superficial and deep masseter) as a positive stimulus for myofascial neurons. Regime C (Figs. 4–5) restricted stimulation to the masseter. Sequential von Frey stimulation (2–60 g) was followed by pinch in CFA experiments, whereas inflammatory soup experiments consisted of von Frey stimulation (2–10 g) followed by repeated massage (4×). Experiments were performed in *Scn10a*-Cre::GCaMP6f mice, and KCl (0.5 M) and capsaicin (100 µM) were applied as positive stimuli to identify myofascial afferents at the end of the experiment.

**Spatial activity maps, response detection, and analysis of fluorescence dynamics**

Spatial activity maps and regions of interest (ROIs) were generated as described previously with minor modifications (16). Activity evoked by repetitive mechanical stimulation was visualized as the pixel-wise standard deviation over time, and stimulus-evoked responses were visualized by subtracting mean pre-stimulus fluorescence from mean fluorescence during stimulation. ROIs were manually selected according to experimental design. In regime A (viral GCaMP6f expression from the masseter), ROIs were selected from neurons responsive to mechanical stimulation. In regime B, neurons responsive to KCl (0.5 M) were separated from neurons responding exclusively to mechanical stimulation to distinguish myofascial from non-myofascial populations. In regime C (*Scn10a*-Cre::GCaMP6f mice), ROIs were selected from neurons responsive to KCl, capsaicin, or both. Relative fluorescence changes were calculated as ΔF/F for each cell. To minimize contamination from out-of-focus tissue and neighboring cells, fluorescence from the surrounding neuropil was subtracted using a custom MATLAB script (16).

Responses were detected from baseline-corrected ΔF/F traces using a custom MATLAB pipeline (16). Baseline fluorescence was defined as the mean signal during the 40 frames preceding stimulus onset. Unless otherwise indicated, responses were defined as events of at least 10% ΔF/F lasting at least 1 frame. All putative positive responses were visually inspected, and only stimulus-locked responses were included in the analysis. This threshold was used to detect, sort, and quantify responses in spatial activity maps using custom Python scripts.

**Quantification and classification of mechanosensory responses**

Only mechanically responsive neurons were included in the analysis. A mechanically responsive neuron was defined as exhibiting at least one stimulus-locked response to mechanical stimulation and lacking frequent spontaneous, non–stimulus-locked activity. Response magnitude was calculated from baseline-corrected ΔF/F traces over the stimulus response window (160 frames; acquisition rate, 5 Hz), and area under the curve (AUC) was computed over the same interval.

To distinguish PEP neurons from Aδ-HTMRs in *Scn10a*-Cre::GCaMP6f experiments, responses to capsaicin and KCl were thresholded at ΔF/F ≥20%, classifying neurons as PEP if capsaicin-responsive and as Aδ-HTMR if responding exclusively to KCl. This higher threshold was used for chemical classification to distinguish robust chemical responses from potential mechanically evoked signals during solution application or injection. For detection of von Frey responses, neuronal activity was additionally required to correlate with the population-average ΔF/F trace from the same mouse (Pearson's $r \geq 0.3$). The population reference trace was generated from neurons exhibiting at least one response to von Frey stimulation. In CFA experiments, correlations were computed across the full von Frey response window within each 200-frame stimulus block, excluding the 40-frame baseline period. In inflammatory soup (IS) experiments, population-level activation events were first identified from the population-average ΔF/F trace using peak detection (scipy.signal.find_peaks, prominence = 0.5) restricted to response-period frames. For each detected peak, a ±10-frame window was defined, and correlations were computed between each neuron's ΔF/F trace and the population-average trace restricted to these event windows. A von Frey response was classified as positive only if it met both the correlation criterion and the ≥10% ΔF/F threshold.

### Application and grouping of manual mechanical stimuli

Manual mechanical stimuli (massage, stretch, pinch) were applied in a standardized manner across animals. Massage stimulation consisted of rhythmic rotational compression of the exposed muscle surface using gentle digital contact during the stimulus window. The finger was placed on the muscle and moved in a controlled circular motion, displacing tissue over the underlying bone without deep indentation. Applied pressure was regulated to avoid skull displacement that could interfere with trigeminal ganglion imaging and produce movement-related artifacts. Massage was applied continuously throughout the response window. Stretch was induced by controlled jaw manipulation, including repeated lateral displacement (5×) and maximal opening (5×). Pinch stimuli consisted of either repeated forceps pinches (10×) or a sustained pinch using an alligator clip; in CFA experiments, standardized pinches (6×) were delivered using an alligator clip. Stimuli were applied with consistent timing and in a standardized sequence of increasing mechanical intensity. Von Frey stimulation consisted of indentations at five distinct locations on the masseter, each repeated twice (10 applications per filament total) with consistent interstimulus intervals.

When multiple closely related mechanical maneuvers were applied within a stimulus class (e.g., massage using index finger or thumb; brushing with or against the grain; different vibration frequencies or amplitudes; prolonged clip pinch and repeated forceps pinch; jaw stretch paradigms), responses were grouped into a single stimulus category for analysis. For each neuron, the response with the largest AUC within that category was used to represent that stimulus class.

### Threshold and recruitment analyses

Von Frey response threshold was defined as the lowest force eliciting a positive response that was followed by a positive response to the next higher-force filament, or, for the highest force tested, a positive response at that force. Only neurons fulfilling this criterion were included in threshold-based recruitment analyses. Cumulative recruitment curves were calculated as the fraction of neurons with thresholds at or below each force. In IS experiments, the median recruitment force was obtained by linear interpolation at 50% recruitment from the empirical cumulative distribution; recruitment curves were additionally fitted with a power function for visualization and nested-model comparison (separate versus combined fits) using an F-test. In CFA experiments, cumulative recruitment curves were fitted with a saturating exponential function constrained to 100% maximal recruitment, and EC50 values were derived from the fitted rate constant.

### Cell-size analysis of chemically identified neurons

For fig. S6, cell-size analysis was performed independently of the functional classification used for von Frey response analysis. Cell size was first measured in Fiji/ImageJ as ROI area in pixels$^2$ from SD images of $Ca^{2+}$ transients evoked by capsaicin or KCl. Neurons were then separated into capsaicin-responsive and KCl–only groups at SD thresholds of 50, 100, and 150 using a custom Python script.

**Behavioral assays**

To determine whether activation of myofascial *Scn10a*$^+$ neurons influences spontaneous behavior, mice were tested in the open field assay. This paradigm quantifies exploratory locomotion and spatial preference in a novel environment. Mice were briefly anesthetized with isoflurane (2% in oxygen) for intramuscular injection of clozapine-N-oxide (CNO; 0.02 mg/kg; 0.025 mg/ml) into the masseter muscle. We previously showed that this low dose does not induce nocifensive behavior or aversive states when delivered systemically (22). After recovery from anesthesia, animals were returned to their home cage for 10 min before being placed in a 42 × 42 cm open field arena for 20 min of exploration under standard ambient lighting conditions. Total distance traveled and center entries were quantified using automated tracking software (TopScan, CleverSys Inc.). Age- and sex-matched heterozygous Scn10a-Cre control mice and Scn10a-Cre::hM3Dq mice received identical CNO injections and were analyzed in parallel.

To validate chemogenetic activation in a peripheral site that reliably evokes overt nocifensive behavior, the same animals were also tested after CNO injection into the glabrous skin of the left hindpaw. Mice were habituated to the experimental room and testing setup before injection, then placed on a wire rack in clear acrylic chambers (10 × 10 × 13 cm) and video recorded from below for 30 min after injection. Each mouse received a 20 µl injection of CNO (0.02 mg/ml) into the glabrous skin of the left hindpaw, corresponding to a dose roughly comparable to that used for intramuscular masseter injection. The first 15 min were analyzed by an experimenter blinded to genotype and separate from the experimenter who performed the injections and recordings. Videos were scored in BORIS, with paw shakes and withdrawals scored as point events and licking or attending of the injected paw scored as state behaviors.

A massage behavioral assay was developed for this study. Six-week-old rats (Long-Evans) were habituated for two weeks prior to testing through repeated handling sessions every other day. During habituation, animals were gradually acclimated to manual massage of the masseter muscle until stimulation was tolerated without overt signs of anxiety. During habituation, animals were handled and acclimated to massage by both the experimenter administering massage during testing and the experimenter responsible for video recording. During testing, massage was applied manually using the index finger to one masseter muscle for up to 30 s. If tolerated for the full duration, stimulation was subsequently applied to the contralateral side; otherwise, sides were switched upon withdrawal. Each side was tested six times per session. Massage tolerance duration was determined in a blinded manner from recorded videos and averaged for each side. Withdrawal behaviors were defined as turning away from the stimulus or actively pushing the experimenter's finger away. Baseline massage tolerance was assessed on test day 1. To induce inflammation, complete Freund's adjuvant (CFA; 100 µl) was injected intramuscularly into one masseter muscle the following day, with injection side balanced across animals. Behavioral testing was repeated 24 h later. The experimenter administering massage was blinded to injection side. Although CFA frequently produced visible swelling that could partially reveal injection side, massage was delivered according to a predefined standardized protocol independent of side to minimize bias.

**In situ hybridization and post hoc molecular identification**

To determine the molecular identity of masseter-innervating neurons, C57BL/6 mice (2–4 months) received intramuscular AAV6.2-Cre-mCherry injections into the masseter. Seven days later, trigeminal ganglia were harvested, cryosectioned (20 µm), and processed using multiple rounds of hybridization chain reaction (HCR; Molecular Instruments), as described previously (12). Probes targeting *Nppb, Sst*, *Tmem233*, *Fxyd2*, *Mrgprd*, *S100b*, *Scn10a*, *Trpm8*, *Trpv1*, *Calca*, and *mCherry* were used to classify mCherry-positive neurons. For post hoc molecular identification following in vivo calcium imaging, trigeminal ganglia were fixed immediately after imaging and processed using HCR v3 (15) with probes targeting *S100b*, *Scn10a*, *Calca*, and *mCherry*. Whole-mount ganglia were imaged using an Olympus FV3000 confocal microscope. Sparse mCherry expression enabled re-identification of imaged neurons between calcium imaging and HCR datasets. Only neurons in which probe signal could be confidently attributed to the corresponding soma were included for molecular classification; ambiguous cases were excluded after cross-validation across probe channels.

**Whole-mount clearing and immunolabeling**

Whole-mount clearing of the masseter and craniofacial tissues was performed with modifications of the KneEZ Clear protocol (38). *Scn10a*-Cre; Ai140 mice were perfused with PBS followed by 4% PFA. Heads were post-fixed, hemisected, and decalcified (10% EDTA, 6 days). Samples underwent graded tetrahydrofuran dehydration and rehydration, followed by refractive index equilibration in EZ View solution. For immunolabeling, tissues were permeabilized, blocked, and incubated with sheep anti-GFP (1:500, 4 days) and donkey anti-sheep Alexa Fluor 647 (1:1000). Samples were cleared in EZ Clear (RI 1.51) and imaged on an Olympus FV3000 confocal microscope as stitched z-stacks. Whole-mount optical clearing of brain and cervical tissue was performed following the LifeCanvas Whole Mouse Head protocol (39). Samples were perfused, post-fixed, decalcified, SHIELD-stabilized, and subjected to organic solvent–based delipidation. Cleared tissue was refractive index–matched in EasyIndex (RI 1.52) and imaged using a SmartSPIM light-sheet microscope.

**Statistical analysis**

Statistical analyses were performed in Python (SciPy) unless otherwise indicated. For behavioral datasets, paired and unpaired nonparametric comparisons were used as appropriate to the experimental design (paired: Wilcoxon signed-rank test; unpaired: Mann–Whitney U test). For von Frey threshold comparisons between independent groups, Mann–Whitney U tests were applied. Differences in categorical response distributions were assessed using pairwise Fisher's exact tests, with Holm correction for multiple comparisons where applicable. For cumulative recruitment analyses, nonlinear regression was used to fit saturating exponential or power functions as specified, and nested-model comparisons (separate versus shared fits) were evaluated using F-tests based on residual sums of squares. All tests were two-sided unless otherwise noted. Data are presented as mean ± s.e.m. unless otherwise indicated. Box plots indicate the median (center line), interquartile range (box), and 1.5× interquartile range (whiskers); individual data points are overlaid. n denotes individual neurons unless otherwise specified, and the number of animals used for each experiment is indicated in the corresponding figure legends. No statistical methods were used to predetermine sample size. Animals were allocated to experimental groups without formal randomization. $P < 0.05$ was considered statistically significant.

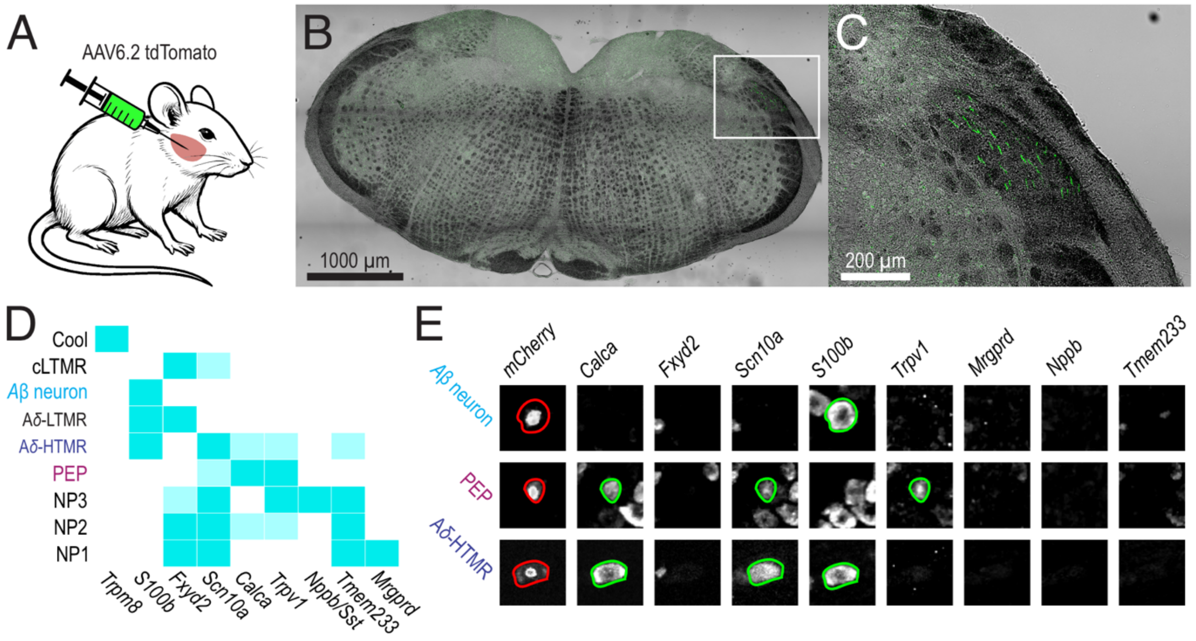


**Fig. S1. Trigeminal masseter afferents and molecular classification scheme.**
(**A**) Schematic of AAV6.2-tdTomato injection into the masseter muscle. (**B**) Brainstem section showing the spinal trigeminal subnucleus caudalis (Sp5c). (**C**) Higher-magnification view of the boxed region in (**B**), showing tdTomato$^{+}$ fibers in the dorsal part of the Sp5c. (**D**) Molecular classification scheme for nine sensory neuron classes. Boxes indicate high (dark) or lower (faint) expression of each marker in the indicated class. (**E**) Examples of mCherry$^{+}$ neurons from the three major classes identified among masseter afferents and the molecular markers they express.

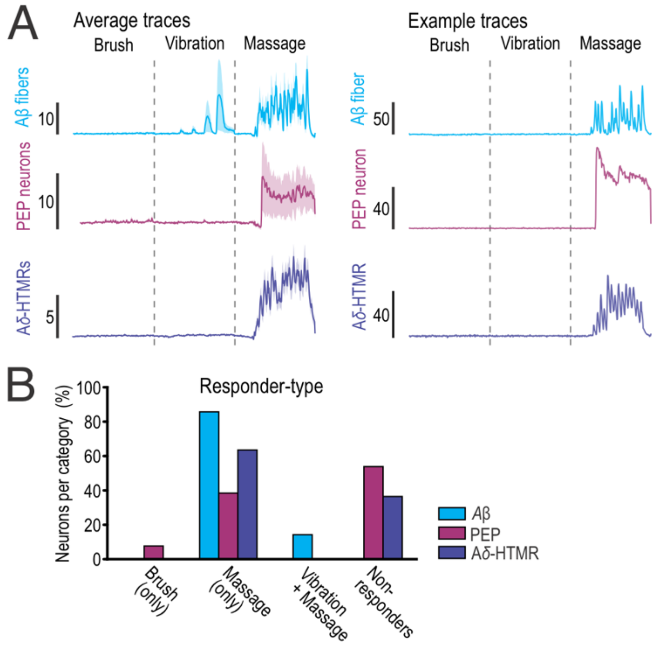


**Fig. S2. Massage recruits masseter afferents through the skin.**
(**A**) Average and example traces of the three major classes shown in Fig. 1G, displaying responses to stimulation through the intact skin. Gentle cutaneous stimuli such as brush and vibration rarely activated masseter afferents, whereas massage recruited neurons in each class. Average traces are shown for all neurons within each class ± s.e.m. $n$ = 7 Aβ neurons, 26 PEP neurons, and 85 Aδ-HTMR neurons from 5 mice. (**B**) Quantification of responder types during cheek stimulation. All Aβ neurons, 63.5% of Aδ-HTMRs, and 46.1% of PEP neurons responded to massage. Among neurons that responded to direct muscle stimulation, 53.8% of PEP neurons and 36.5% of Aδ-HTMRs did not respond to any stimulus applied through the skin, whereas all Aβ neurons responded to at least one cheek stimulus.

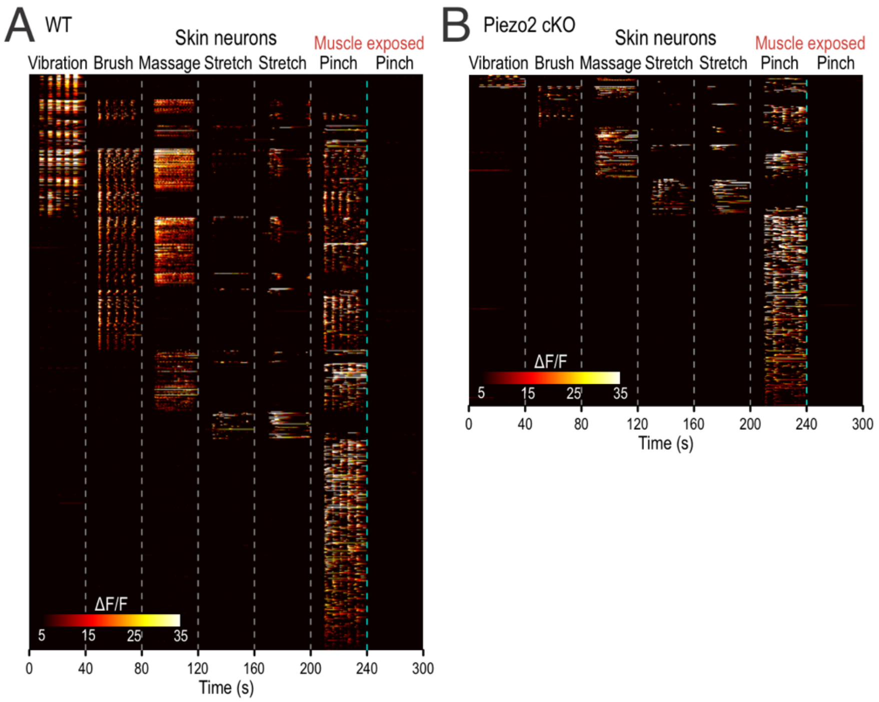


**Fig. S3. Skin neuron responses in WT and Piezo2 cKO mice.**
(**A**) Population heatmaps of skin neurons in WT mice from Fig. 2C. Neurons were classified as cutaneous if they responded to skin stimulation but not to subsequent direct stimulation of the exposed masseter after skin removal. These neurons showed the expected cutaneous response profile, with strong responses to brush and vibration. Massage and jaw stretch were also applied before skin removal. For stretch, the first stimulus consisted of repeated jaw stretches and the second of a prolonged jaw stretch. Pinch of the exposed masseter, the strongest muscle stimulus, is shown to illustrate that these neurons responded only to cutaneous stimulation. *n* = 958 neurons from 3 mice. (**B**) Same analysis in Piezo2 cKO mice from Fig. 2D. *n* = 567 neurons from 4 mice.

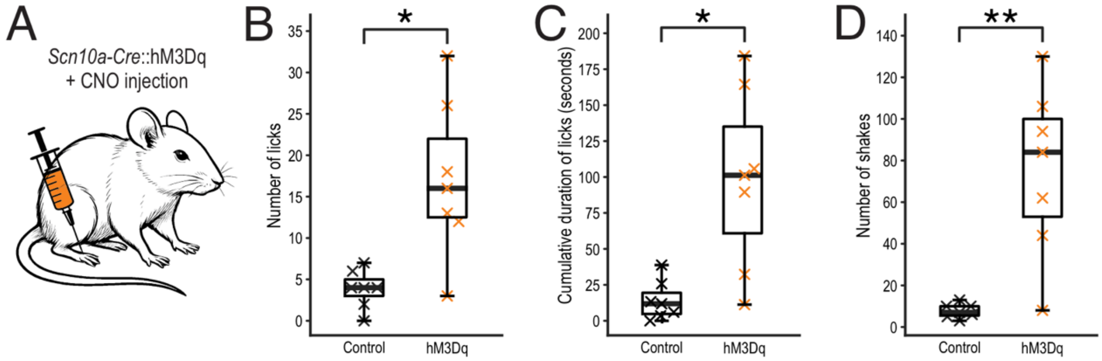


**Fig. S4. Chemogenetic activation of *Scn10a*⁺ neurons in the hindpaw.**
(**A**) Schematic of CNO injection into the hindpaw of *Scn10a*-Cre::hM3Dq mice. WT and hM3Dq mice received identical CNO dosing. The same animals were used as in Fig. 3E–H. *n* = 7 mice per group. (**B**) Quantification of licks directed toward the injected hindpaw. Licking was significantly increased in hM3Dq mice (WT: 3.86; hM3Dq: 17.14; Mann–Whitney U test, *p* = 0.0148). (**C**) Cumulative licking duration directed toward the injected hindpaw was also increased in hM3Dq mice (WT: 14.17 s; hM3Dq: 98.39 s; Mann–Whitney U test, p = 0.011). (**D**) Hindpaw shakes were likewise increased in hM3Dq mice (WT: 7.71; hM3Dq: 75.43; Mann–Whitney U test, p = 0.007).

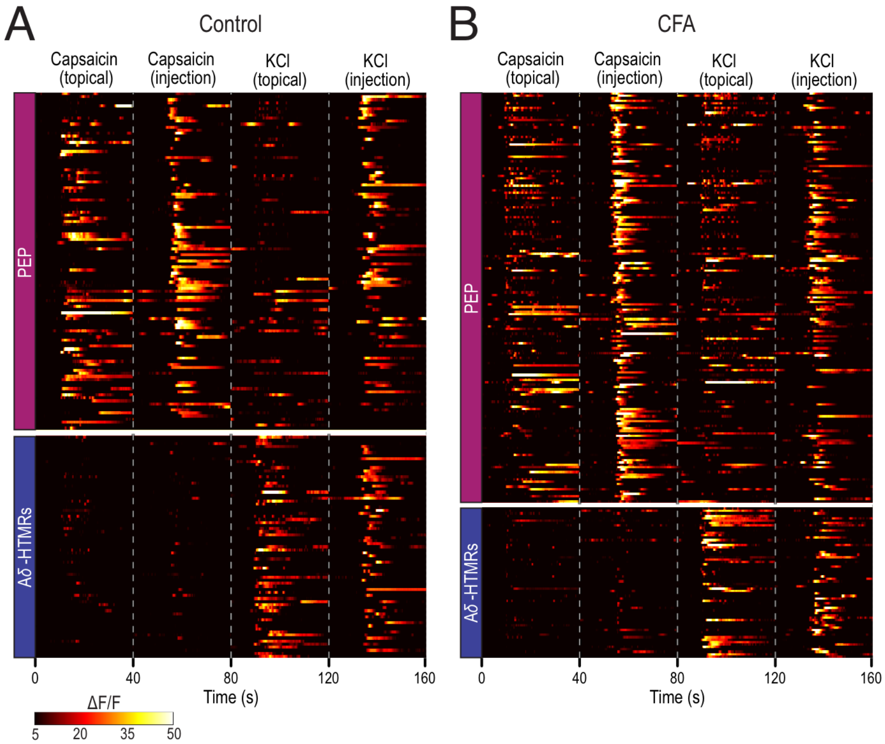


**Fig. S5. Functional separation of PEP and Aδ-HTMRs by capsaicin and KCl responses.**
(**A**) Population heatmaps showing responses to topical application and intramuscular injection of capsaicin and high KCl in *Scn10a*-Cre::GCaMP6f mice under control conditions. Neurons showing robust capsaicin responses were assigned to the PEP class, whereas neurons showing robust KCl responses but lacking capsaicin responses were assigned to the Aδ-HTMR class. PEP neurons: *n* = 110; Aδ-HTMRs: *n* = 73 neurons from 5 mice. (**B**) Same analysis in *Scn10a*-Cre::GCaMP6f mice under CFA conditions. PEP neurons: *n* = 169; Aδ-HTMRs: *n* = 63 neurons from 6 mice.

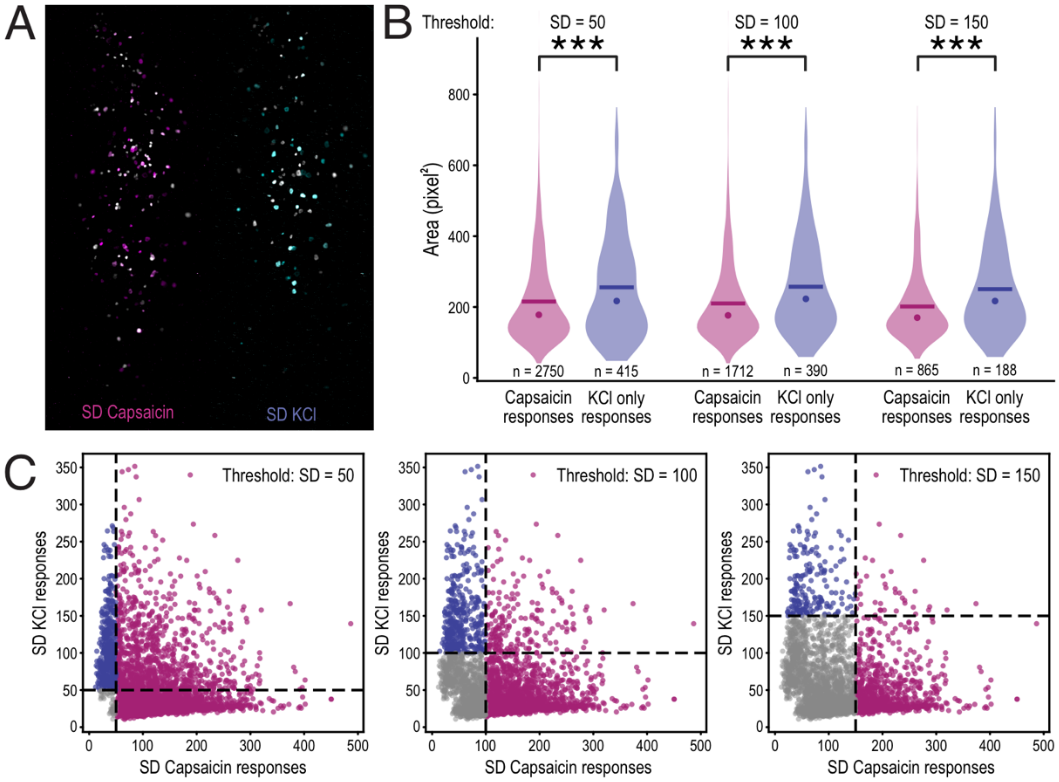


**Fig. S6. Cell-size distribution of capsaicin-responsive and KCl–only neurons.**
(**A**) Representative standard-deviation (SD) images of $Ca^{2+}$ transients during topical application and intramuscular injection of capsaicin (topical: gray; intramuscular: magenta, left) or KCl (topical: gray; intramuscular: cyan, right) in *Scn10a*-Cre::GCaMP6f mice. (**B**) Violin plots of cell size (pixels²) for capsaicin-responsive neurons and high KCl–only neurons across three SD thresholds (Mann–Whitney U test: SD = 50, $p = 8.7 \times 10^{-7}$; SD = 100, $p = 1.1 \times 10^{-11}$; SD = 150, $p = 8.7 \times 10^{-7}$). Means are indicated by lines and medians by dots. The number of neurons is indicated for each threshold; data were pooled across control and CFA experiments from 10 mice. (C) Plots showing neurons selected at each threshold (blue: KCl only; magenta: capsaicin-responsive; gray: excluded). Excluded neurons: SD = 50, $n = 51$; SD = 100, $n = 1114$; SD = 150, $n = 2163$.

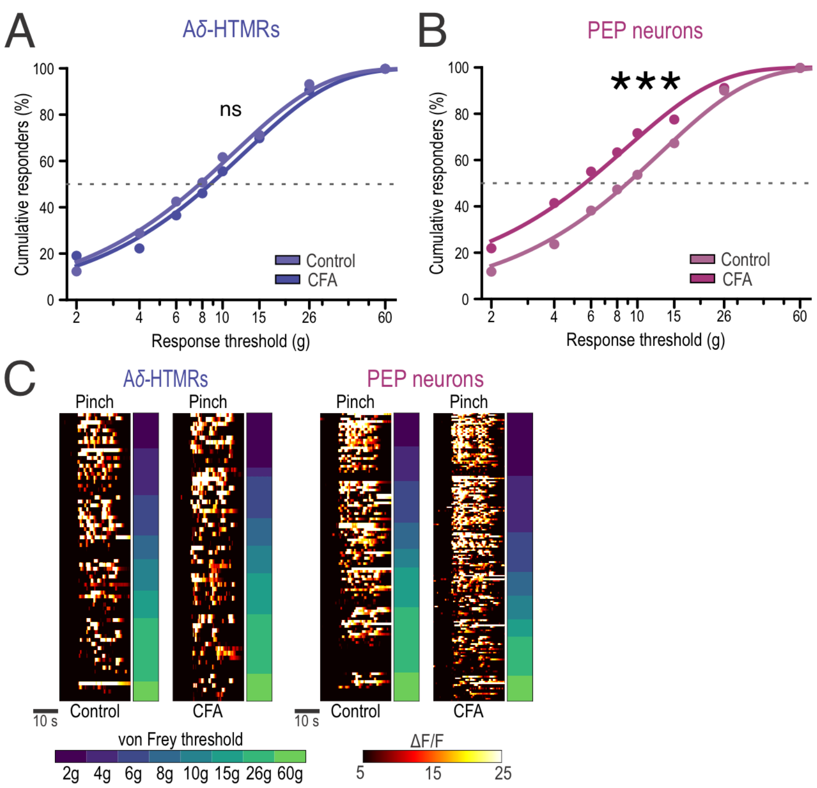


**Fig. S7. Inflammation-dependent recruitment curves and pinch responses.**

(**A**) Force–response curves of Aδ-HTMRs under control and CFA conditions. Mechanical sensitivity did not differ significantly between control and CFA conditions (Control EC50 = 7.76 ± 0.46 g; CFA EC50 = 8.74 ± 0.61 g; nonlinear regression comparison, F-test, $p = 0.111$). Control: $n = 73$ neurons from 5 mice; CFA: $n = 63$ neurons from 6 mice. (**B**) Same analysis for PEP neurons. CFA, PEP neurons were significantly more sensitive (ΔEC50 = 2.77 g; control EC50 = 8.95 ± 0.46 g; CFA EC50 = 6.18 ± 0.51 g; nonlinear regression comparison, F-test, $p = 5.34 \times 10^{-6}$). Control: $n = 110$ neurons from 5 mice; CFA: $n = 169$ neurons from 6 mice. (**C**) Left, heatmap of pinch responses in von Frey-responsive Aδ-HTMRs under control and CFA conditions, shown alongside color-coded von Frey thresholds. Right, same analysis for PEP neurons. In both populations, pinch responses spanned the full von Frey threshold range under control and CFA conditions.

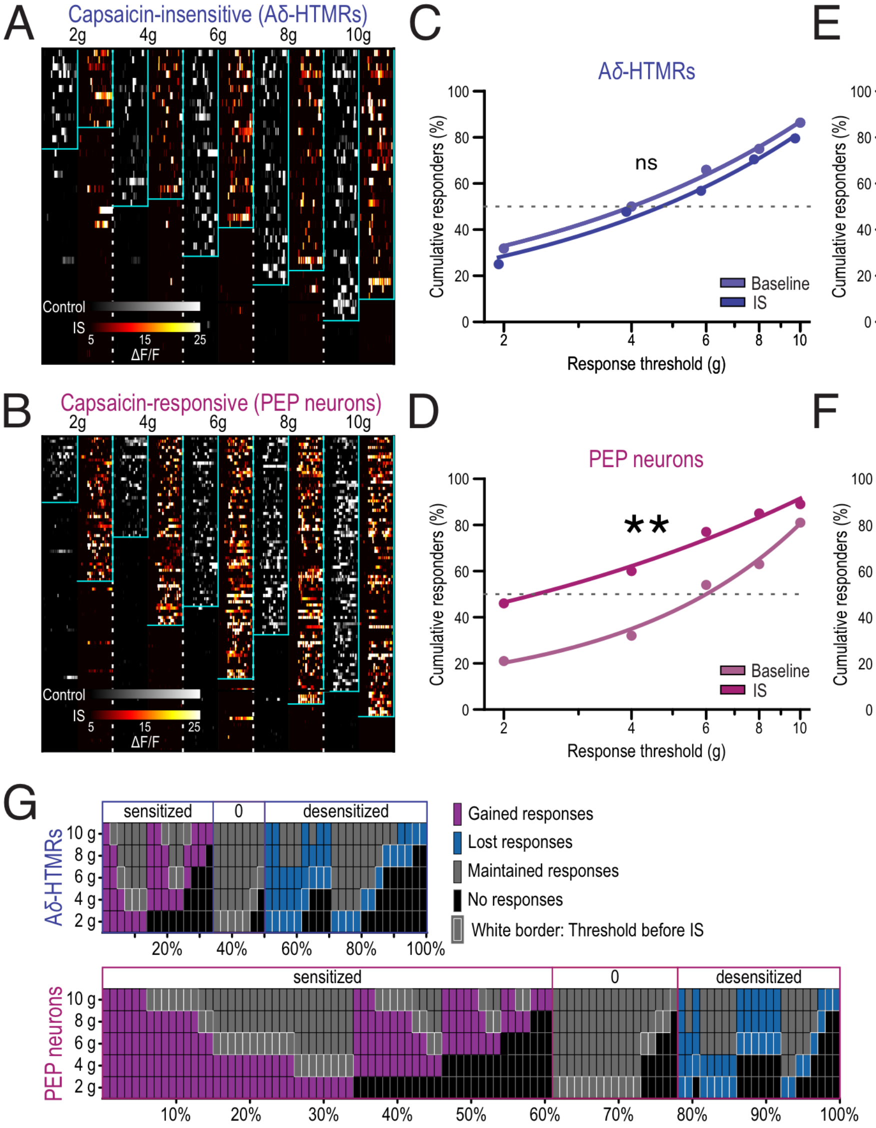


**Fig. S8. IS heatmaps and inflammation-dependent recruitment curves.**
(**A**) Longitudinal in vivo $Ca^{2+}$ imaging before and 4 min after acute inflammatory soup (IS) injection into the masseter muscle of *Scn10a*-Cre::GCaMP6f mice. Population heatmaps show alternating von Frey responses (2–10 g) for capsaicin-insensitive Aδ-HTMRs (gray, control; fire, IS). Control and IS responses were sorted independently; each stimulus block was 40 s long. $n$ = 44 neurons from 8 mice. (**B**) Same analysis for capsaicin-sensitive PEP neurons. $n$ = 100 neurons from 8 mice. (**C**) Force–response curves of Aδ-HTMRs before and after IS. Mechanical sensitivity did not differ significantly after IS (50% recruitment: control = 4.0 g, IS = 4.5 g; F-test, $p$ = 0.118). (**D**) Same analysis for PEP neurons. IS significantly increased PEP neuron sensitivity (50% recruitment: control = 5.64 g, IS = 2.57 g; Δ = 3.07 g; F-test, $p$ = 0.003). (**E**) Force–response curves comparing Aδ-HTMRs and PEP neurons under baseline conditions. Aδ-HTMRs were more mechanically sensitive than PEP neurons (50% recruitment: Aδ = 4.0 g, PEP = 5.64 g; Δ = 1.64 g; F-test, $p$ = 0.0153). (**F**) Same comparison after IS. IS increased PEP neuron sensitivity, reversing the subtype relationship observed under baseline conditions (50% recruitment: PEP = 2.57 g, Aδ = 4.5 g; Δ = 1.93 g; F-test, $p$ = 0.0098). (**G**) Single-neuron sensitivity-shift analysis across 2 g force steps. Each column represents one neuron. Gained responses after IS are shown in magenta, lost responses in blue, maintained responses in gray, and no response in black; white borders indicate the pre-IS response threshold. Aδ-HTMRs showed mixed changes biased toward response loss, whereas PEP neurons showed a shift toward increased sensitivity.

**Movie S1. Rat massage assay under control conditions.**
Short video example from the control session shown in Fig. 5. The rat received cheek massage on both sides on test day 1 and tolerated massage on both sides for approximately equal durations. This behavior was observed repeatedly during the session.

**Movie S2. Rat massage assay after CFA-induced inflammation.**
Short video example from the same rat shown in Movie S1, recorded 24 h after unilateral CFA injection. The rat tolerated massage of the left, non-injected side for the maximum duration. When massage was applied to the inflamed, CFA-injected right side, the rat withdrew and batted at the experimenter's finger after a short latency. This behavior was observed repeatedly during the session.
